# Systematic detection of brain protein-coding genes under positive selection during primate evolution and their roles in cognition

**DOI:** 10.1101/658658

**Authors:** Guillaume Dumas, Simon Malesys, Thomas Bourgeron

## Abstract

The human brain differs from that of other primates, but the genetic basis of these differences remains unclear. We investigated the evolutionary pressures acting on almost all human protein-coding genes (*N*=11,667; 1:1 orthologs in primates) based on their divergence from those of early hominins, such as Neanderthals, and non-human primates. We confirm that genes encoding brain-related proteins are among the most strongly conserved protein-coding genes in the human genome. Combining our evolutionary pressure metrics for the protein-coding genome with recent datasets, we found that this conservation applied to genes functionally associated with the synapse and expressed in brain structures such as the prefrontal cortex and the cerebellum. Conversely, several genes presenting signatures commonly associated with positive selection appear as causing brain diseases or conditions, such as micro/macrocephaly, Joubert syndrome, dyslexia, and autism. Among those, a number of DNA damage response genes associated with microcephaly in humans such as *BRCA1, NHEJ1, TOP3A*, and *RNF168* show strong signs of positive selection and might have played a role in human brain size expansion during primate evolution. We also showed that cerebellum granule neurons express a set of genes also presenting signatures of positive selection and that may have contributed to the emergence of fine motor skills and social cognition in humans. This resource is available online and can be used to estimate evolutionary constraints acting on a set of genes and to explore their relative contributions to human traits.

## Introduction

Modern humans (*Homo sapiens*) can perform complex cognitive tasks well and communicate with their peers (Dunbar and Shultz 2017). Anatomic differences between the brains of humans and other primates are well documented (e.g., cortex size, prefrontal white matter thickness, lateralization), but how the human brain evolved remains a matter of debate (Varki et al. 2008). A recent study of endocranial casts of *Homo sapiens* fossils indicates that brain size in early *Homo sapiens*, 300,000 years ago, was already within the range of that in present-day humans (Neubauer et al. 2018). However, brain shape, evolved more gradually within the *Homo sapiens* lineage, reaching its current form between about 100,000 and 35,000 years ago. It has also been suggested that the enlargement of the prefrontal cortex relative to the motor cortex in humans is mirrored in the cerebellum by an enlargement of the regions of the cerebellum connected to the prefrontal cortex (Balsters et al. 2010). These anatomic processes of tandem evolution in the brain paralleled the emergence of motor and cognitive abilities, such as bipedalism, planning, language, and social awareness, which are mainly well developed in humans.

Genetic differences in primates undoubtedly contributed to these brain and cognitive differences, but the genes or variants involved remain largely unknown. Indeed, demonstrating that a genetic variant is adaptive requires strong evidence at both the genetic and functional levels. Only a few genes have been shown to be human-specific. They include *SRGAP2C* (Charrier et al. 2012), *ARHGAP11B* (Florio et al. 2015), and *NOTCH2NLA* (Suzuki et al. 2018), which emerged through recent gene duplication in the *Homo* lineage (Dennis et al. 2017). The expression of these human-specific genes in the mouse brain expand cortical neurogenesis (Florio et al. 2015; Suzuki et al. 2018; Nuttle et al. 2016; Dennis et al. 2012). Several genes involved in brain function display accelerated coding region evolution in humans. For example, *FOXP2* has been associated with verbal apraxia and *ASPM* with microcephaly (Enard et al. 2002; Montgomery et al. 2014). Functional studies have also shown that mice carrying a “humanized” version of *FOXP2* display qualitative changes in ultrasonic vocalization (Enard et al. 2009). However, these reports targeting only specific genes sometimes provide contradictory results (Atkinson et al. 2018). Other studies have reported sequence conservation to be stronger in the protein-coding genes of the brain than in those of other tissues (Miyata et al. 1994; Wang et al. 2006; Tuller et al. 2008), suggesting that the primary substrate of evolution in the brain is regulatory changes in gene expression (King and Wilson 1975; Pollard et al. 2006; Changeux 2017) and splicing (Calarco et al. 2007). In addition, several recent studies have recently explored the genes subjected to the highest degrees of constraint during primate evolution or in human populations, to improve estimations of the pathogenicity of variants identified in patients with genetic disorders (Sundaram et al. 2018; Havrilla et al. 2019). By contrast, fewer studies have systematically detected genes that have diverged during primate evolution (Dorus et al. 2004; Huang et al. 2013; Nielsen et al. 2005).

We describe here an exhaustive screening of all protein-coding genes for conservation and divergence from the common primate ancestor, making use of rich datasets of brain single-cell transcriptomics, proteomics, and imaging to investigate the relationships between these genes and brain structure, function, and diseases.

## Results

### Strong conservation of brain protein-coding genes

We first compared the sequences of modern humans, archaic humans, and other primates to those of their common primate ancestor (inferred from the Compara 6-way primate Enredo, Pecan, Ortheus multiple alignments (Paten et al. 2008)), to extract a measurement of evolution for 11,667 of the 1:1 orthologs across primates, selected from the 17,808 protein-coding genes in the modern human genome (Fig. 1A, see also Fig. S1 and S2; Kapheim et al. 2015). This resource is available online from https://genevo.pasteur.fr/. Our measurement is derived from one of the most widely used and reliable measurements of evolutionary pressure on protein-coding regions, the *d*_N_/*d*_S_ ratio (Yang and Bielawski 2000), also called ω. This measurement compares the rates of non-synonymous and synonymous mutations of coding sequences. If there are more non-synonymous mutations than expected, there is signs of positive selection, if fewer, there is signs of selective constraint. We first estimated *d*_N_ and *d*_S_ for all 1:1 orthologous genes, because the evolutionary constraints on duplicated genes are relaxed (O’Toole et al. 2018) (note: only the Y chromosome was excluded from these analyses). We then adjusted the *d*N/*d*S ratio for biases induced by variations of mutations rate with the GC content of codons. Finally, we renormalized the values obtained for each taxon across the whole genome. The final ω_GC12_ obtained took the form of Z-score corrected for GC content that quantified the unbiased divergence of genes relative to the ancestral primate genome (Kapheim et al. 2015). High positive ω_GC12_ indicates a genetic signature commonly, but not exclusively, associated with positive evolutionary selection; at contrary negative ωGC12 reflects selective constraint.

**Figure 1.**
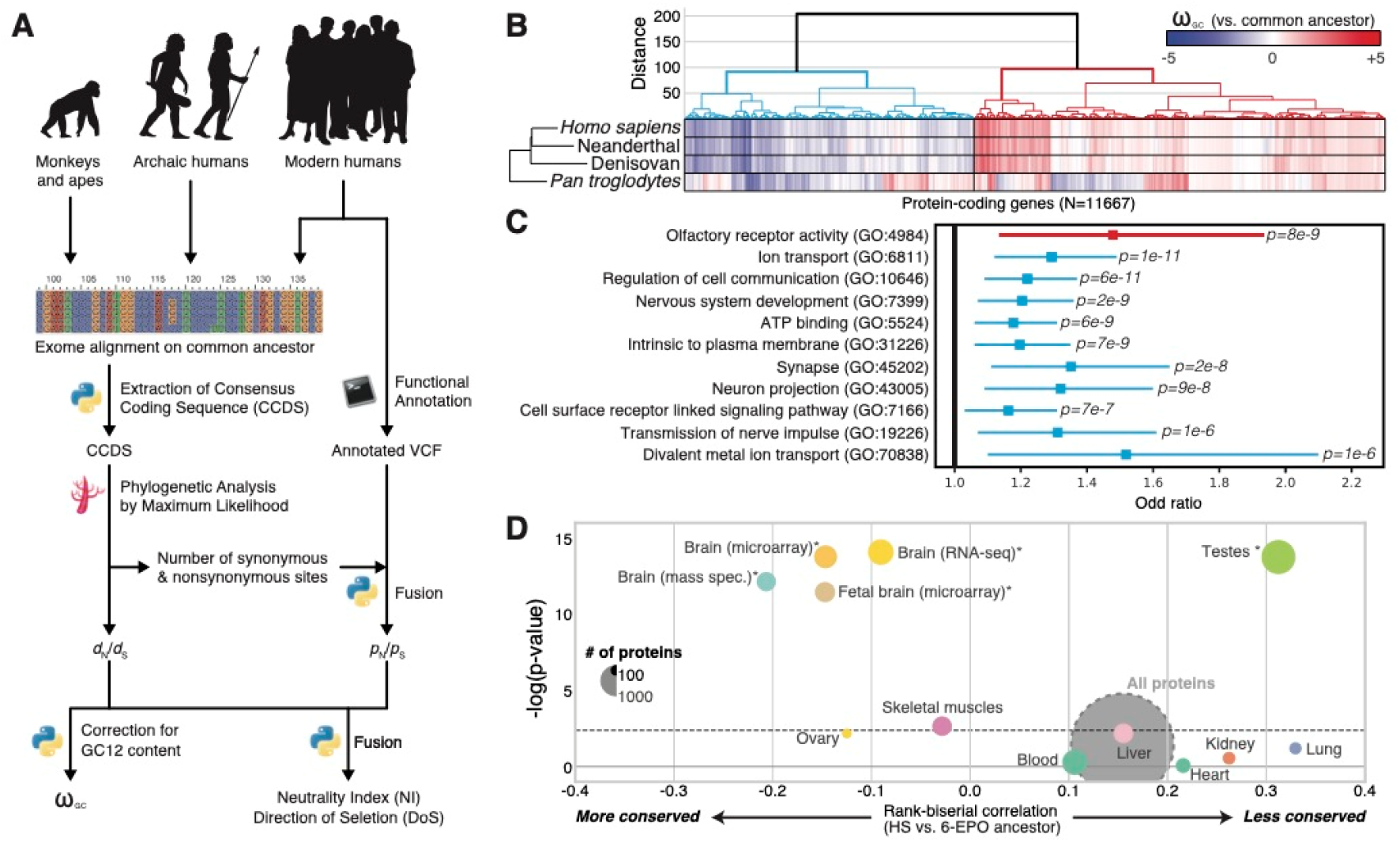
Evolution of protein-coding genes across tissues and biological functions. (A) Analysis pipeline for the extraction of ω_GC12_, a corrected and normalized measurement of the evolution of protein-coding genes that behaves like a Z-score and takes into account the GC content of codons. (B) Hierarchical clustering, based on ω_GC12_, across all protein-coding genes (1:1 orthologs in hominins with medium coverage; See Table S1). (C) Gene Ontology (GO) enrichments for the red and blue clusters in panel b (See Table S2 for all GO terms). Horizontal lines indicate 95% confidence intervals. (D) Funnel plot summarizing the evolution of protein-coding genes specifically expressed in different tissues of the human body (Table S3). Horizontal and vertical axes indicate respectively the effect size and the statistical significance. Circle size indicates the number of proteins in the set. The dashed horizontal line indicates the threshold for significance after Bonferroni correction. Stars indicate the set of genes for which statistical significance was achieved in multiple comparisons after correction, with a bootstrap taking GC12 content and coding sequence length into account. HS: *Homo sapiens;* 6-EPO ancestor: the reconstructed ancestral genome of primates based on alignments of *Homo sapiens*, chimpanzee, gorilla, orangutan, rhesus macaque, and marmoset genomes.

Using the ω_GC12_ for all protein-coding genes in *Homo sapiens*, Denisovans, Neanderthals, and *Pan troglodytes*, we identified two distinct clusters in hominins (Fig. 1B and Table S1): one containing “positively selected” genes (PSG), enriched in olfactory genes (OR=1.48, *p*=8.4×10^-9^), and one with genes under “selective constraint” (SCG), enriched in brain-related biological functions (Fig. 1C and Table S2). This second cluster revealed particularly strong conservation of genes encoding proteins involved in nervous system development (OR=1.2, *p*=2.4×10^-9^) and synaptic transmission (OR=1.35, *p*=1.7×10^-8^).

We investigated the possible enrichment of specific tissues in PSG and SCG by analyzing RNA-seq (Illumina Bodymap2 and GTEx), microarray, and proteomics datasets (Methods). For expression data, despite virtually no gene is expressed only in one tissue, we calculated a tissue specificity score for each genes by normalizing their profile across tissues (see Fig. S3 for more details). The results confirmed a higher degree of conservation for protein-coding genes more specifically expressed in the brain (Wilcoxon rank correlation rc=-0.1, *p*=4.1×10^-12^, bootstrap corrected for gene length and GC content) than for those expressed elsewhere in the body, with the greatest divergence observed for genes expressed in the testis (Wilcoxon rc=0.3, *p*=7.8×10^-11^, bootstrap corrected for gene length and GC content; Fig. 1D, see also Table S3 and Fig. S4 for a replication with GTEx data). This conservation of brain protein-coding genes was replicated with two other datasets (MicroArray: Wilcoxon OR=-0.18, *p*=1.8×10^-12^; mass spectrometry: Wilcoxon rc=-0.21, *p*=1.55×10^-9^; bootstrap corrected for gene length and GC content).

### Conservation of protein-coding genes relating to nervous system substructure and neuronal functions

We then used microarray (Su et al. 2004) and RNA-seq (The GTEx Consortium 2015) data to investigate the evolutionary pressures acting on different regions of the central nervous system. Three central nervous system substructures appeared to have evolved under the highest level of purifying selection at the protein sequence level (ω_GC12_<2): (i) the cerebellum (Wilcoxon rc=-0.29, *p*=5.5×10^-6^, Bonferroni corrected) and the cerebellar peduncle (Wilcoxon rc=-0.11, *p*=3.2×10^-4^, bootstrap corrected for gene length and GC content), (ii) the amygdala (Wilcoxon rc=-0.11, *p*=4.1×10^-6^, bootstrap corrected for gene length and GC content), and, (iii) the prefrontal cortex (Wilcoxon rc=-0.1, *p*=5.7×10^-10^, bootstrap corrected for gene length and GC content; Fig. 2A, see also Table S3). Indeed, it has been suggested that the prefrontal cortex is one of the most divergent brain structure in human evolution (Schoenemann et al. 2005), this diversity being associated with high-level cognitive function (Frith and Dolan 1996). Only one brain structure was expressing more PSG than expected: the superior cervical ganglion (Wilcoxon rc=0.22, *p*=1×10^-6^, bootstrap corrected for gene length and GC content). This structure provides sympathetic innervation to many organs and is associated with the archaic functions of the fight-or-flight response. The PSG expressed in the superior cervical ganglion include *CARF*, which was found to be specifically divergent in the genus *Homo*. This gene encodes a calcium-responsive transcription factor that regulates the neuronal activity-dependent expression of *BDNF* (Tao et al. 2002) and a set of singing-induced genes in the song nuclei of the zebra finch, a songbird capable of vocal learning (Whitney et al. 2014). This gene had a raw *d_N_/d_S_* of 2.44 (7 non-synonymous vs. 1 synonymous mutation in *Homo sapiens* compared to the common primate ancestor) and was found to be one of the PSG with the higher *d*_N_/*d*_S_ value expressed in the human brain.

**Figure 2.**
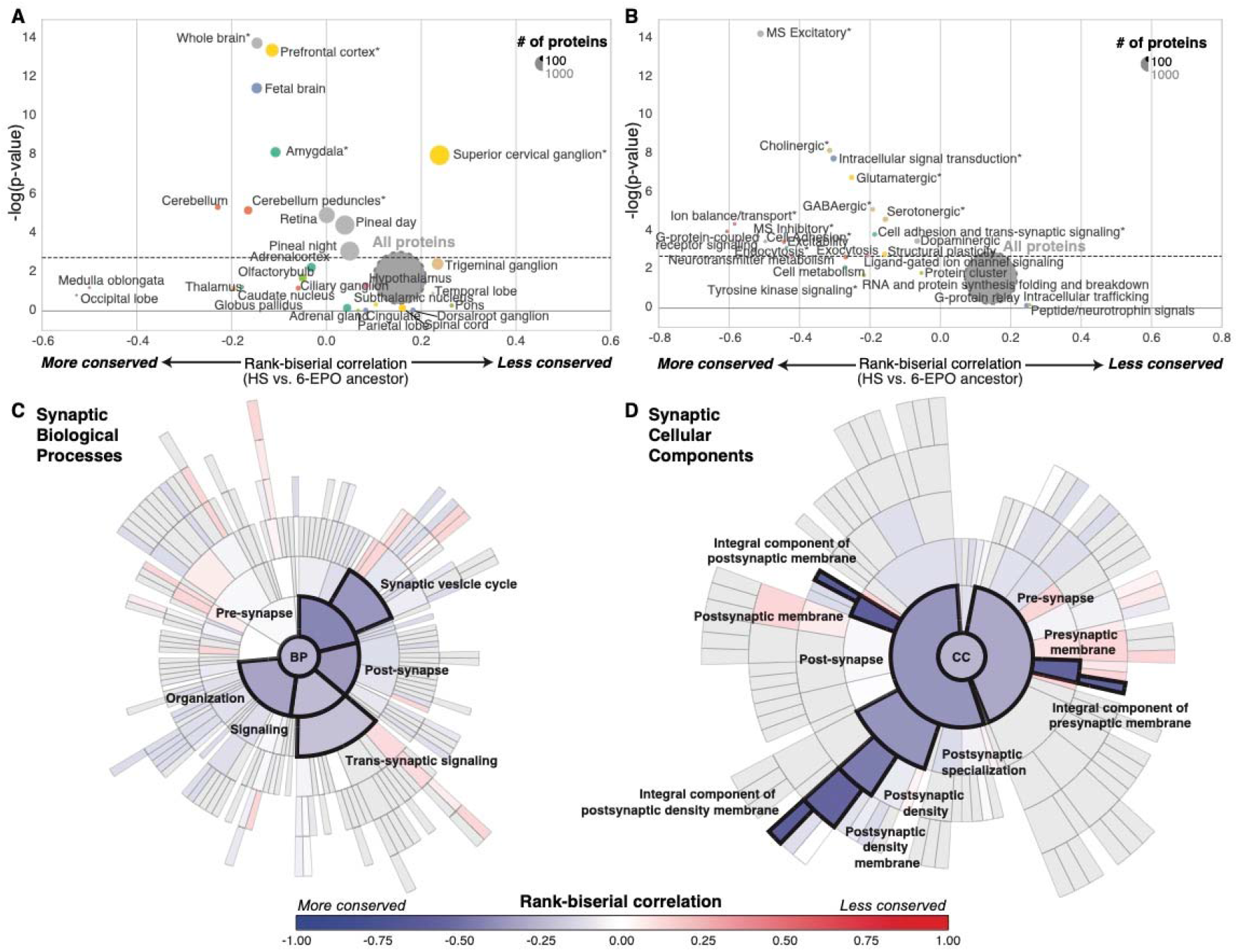
Evolution of brain-related protein-coding genes. (A,B) Funnel plots summarizing the evolution of protein-coding genes specifically expressed in (A) brain substructures and (B) synaptic functions; the dashed horizontal line indicates the threshold for significance after Bonferroni correction. Stars indicate sets of genes for which statistical significance was achieved for multiple comparisons with bootstrap correction; (C, D) SynGO sunburst plots showing nested statistically conserved (blue) biological processes and cellular components of the synapse. The circle in the center represents the root node, with the hierarchy moving outward from the center. A segment of the inner circle bears a hierarchical relationship to those segments of the outer circle which lie within the angular sweep of the parent segment.

We then investigated the possible enrichment of PSG and SCG in brain-specific Gene Ontology terms. All pathways displayed high overall levels of conservation, but genes encoding proteins involved in glutamatergic and GABAergic neurotransmission were generally more conserved (Wilcoxon rc=-0.25; *p*=9.8×10^-6^, Bonferroni corrected) than those encoding proteins involved in dopamine and peptide neurotransmission and intracellular trafficking (Fig. 2B, see also Table S3). The recently released ontology of the synapse provided by the SynGO consortium (http://syngoportal.org) was incorporated into this analysis, not only confirming the globally strong conservation of the synapse but also revealing its close relationship to trans-synaptic signaling processes (Wilcoxon rc=-0.21, p=4.5×10^-5^, Bonferroni corrected) and to postsynaptic (rc=-0.56, p=6.3×10^-8^, Bonferroni corrected) and presynaptic membranes (Wilcoxon: rc=-0.56, *p*=7×10^-8^, Bonferroni corrected; Fig. 2C,D).

### Positively selected genes and their correlation with brain expression and function

We focused on the genes situated at the extremes of the ω_GC12_ distribution (>2SD; Fig. 3A; Table S4) and those fixed in the modern *Homo sapiens* population (neutrality index<1), to ensure that we analyzed PSG with signs of strong positive selection. Only 139 of these 352 highly PSG were brain-related (impoverishment for brain genes, Fisher’s exact test OR=0.66, *p*=1×10^-4^), listed as synaptic genes (Ruano et al. 2010; Lips et al. 2012), specifically expressed in the brain (+2SD for specific expression) or related to a brain disease (extracted systematically from Online Mendelian Inheritance in Man - OMIM: https://www.omim.org and Human Phenotype Ontology - HPO: https://hpo.jax.org/app/). For comparison, we also extracted the 427 SCG under very strong selective constraint, 299 of which were related to the brain categories listed above (enrichment for brain genes, Fisher’s exact test OR=1.26, *p*=0.0032).

**Figure 3.**
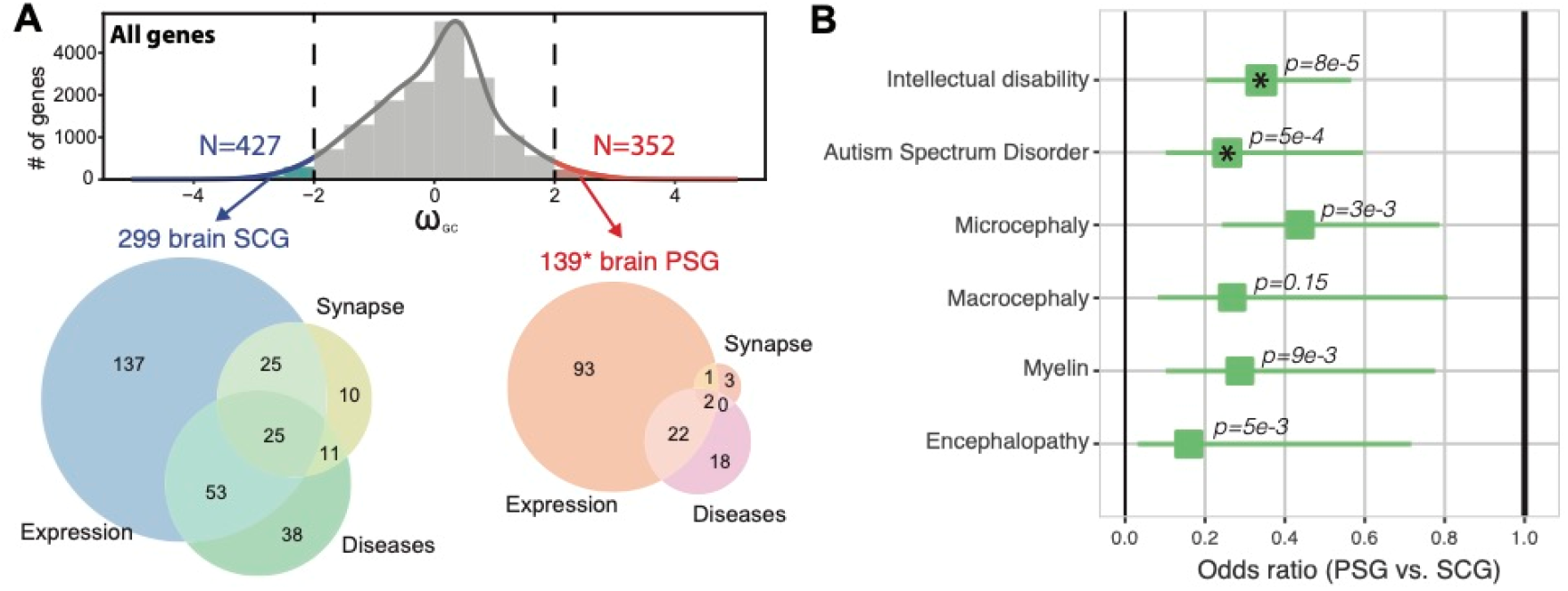
Brain protein-coding genes and human diseases. (A) Distribution of ωGC12 and Venn diagrams describing SCG and PSG situated at the extremes of the ω_GC12_ distribution (>2SD) specifically expressed in the brain (genes with specificity Z-score > 2 in any brain related tissues of Fig. 1d and Fig. 2a), related to the synapse, or brain diseases (Table S4). * addition of 4 genes (*FARSB, KRT14, NPHS1, RSPH1*) containing homo sapiens specific mutations predicted as deleterious (CADD>15). (B) Odds ratios for protein-coding gene sets related to brain diseases (Fisher’s exact test; Asterisks indicate *p*-values significant after Bonferroni correction; horizontal lines indicate 95% confidence intervals).

Using these 427 SCG and 352 PSG, we first used the Brainspan data available from the specific expression analysis (SEA) to confirm that the population of genes expressed in the cerebellum and the cortex was enriched in SCG (Fig. S5). Despite this conservation, based on the adult Allen Brain atlas, we identified a cluster of brain subregions (within the hypothalamus, cerebral nuclei, and cerebellum), more specifically expressing PSG (Fig. S6). Analyses of the human cerebral cortex single cell RNA-seq (Fig. 4A; Table S5; Nowakowski et al. 2017) also revealed an excess of PSG expressed in the choroid plexus — which primary function is to produce cerebrospinal fluid —, in the medial ganglionic eminence (MGE-div) — implicated in the production of GABAergic interneurons and their migration to neocortex during development (Brazel et al. 2003) —, and the radial glial cells (RG).

**Figure 4.**
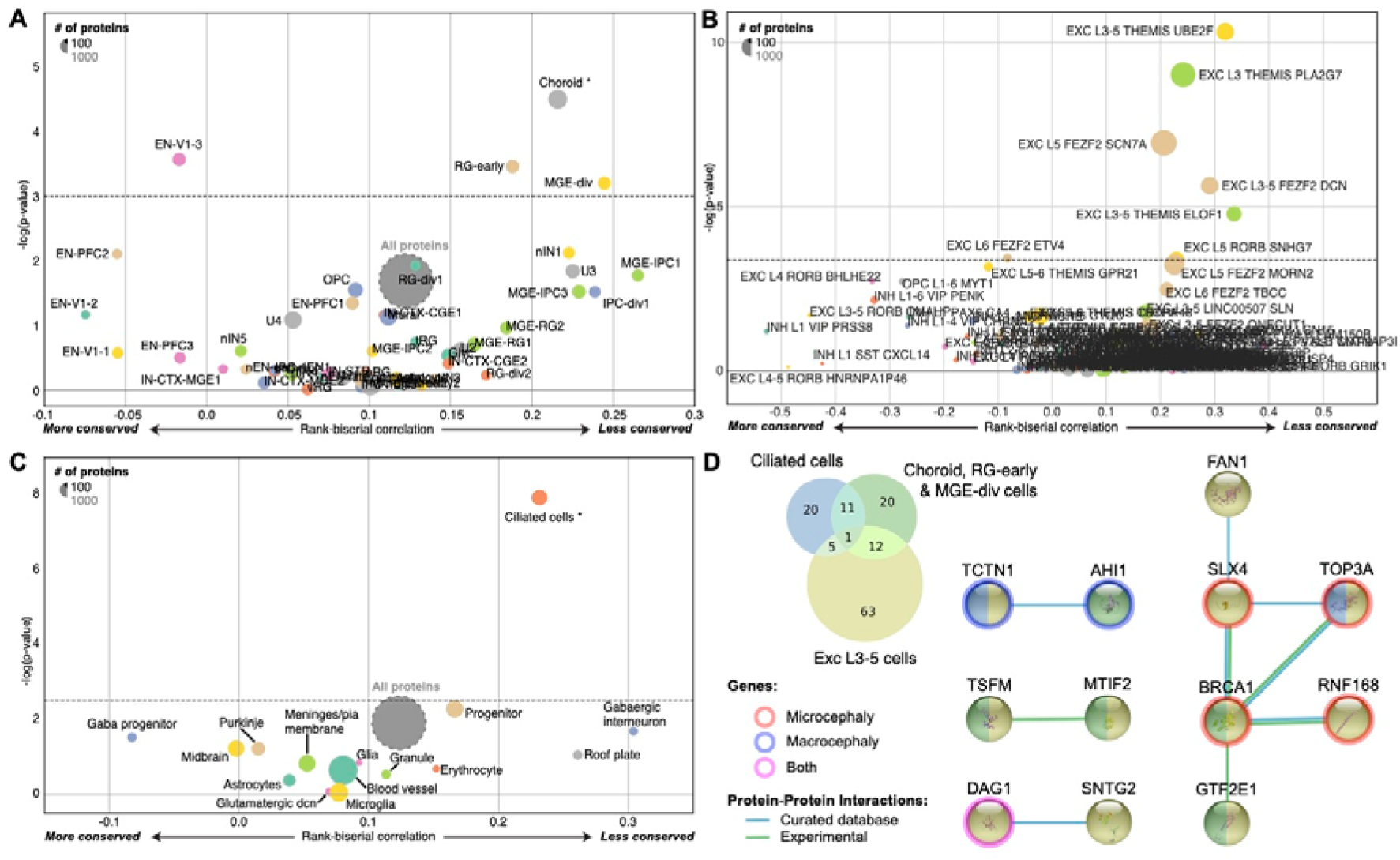
Evolution of protein-coding genes expressed in different cell types. (A, B, C) Funnel plots summarizing the evolution of protein-coding genes specifically expressed in different cell types within (A) the human cerebral cortex (Table S6; Nowakowski et al. 2017), (B) human cortical layers (Table S6; Hodge et al. 2019; Tasic et al. 2018) and (C) the mouse cerebellum (Table S6; Carter et al. 2018). (D) Venn diagram of the PSG expressed specifically in those cell types, with the corresponding Protein-Protein Interaction network (StringDB; Jensen et al. 2009) and their annotated association with micro- and macrocephaly (HPO; Köhler et al. 2019). Abbreviations: EN-V1: primary visual cortex neurons; RG-early: radial glia early cortical progenitors; MGE-div: medial ganglionic eminence dividing cells; Exc: excitatory; l3-5: layers 3-5; Themis, ube2f, pla2g7, etc are cell type markers.

Using a second RNA-seq data set from the human cortex (Hodge et al. 2019; Tasic et al. 2018), we identified 5 cell types, all from layer 3 or 5, expressing PSG more than expected using a stringent Bonferroni and bootstrap correction for gene length and GC content (Fig. 4B; Table S5). Among them, two groups of excitatory neurons — THEMIS PLA2G7 and FEZF2 SCN7A — express several PSG involved in DNA damage response (Arcas et al. 2014) and mutated in patients with microcephaly such as *BRCA1, NHEJ1, RNF168*) and *TOP3A*.

We investigated organoid and human cortex datasets that previously revealed 7 clusters of cells (Camp et al. 2015). Overall the marker genes of these clusters are on average strongly constraint compared to the rest of the genome (Fig. S7). Some PSG are however expressed in these cells, such as *CDC25C, FRMD4B, NHSL1, NUSAP1*, and *PLEKHA5*.

In single-cell transcriptomic studies of the mouse cerebellum (Carter et al. 2018), we found that cells expressing cilium marker genes, such as the dynein light chain roadblock-type 2 (*DYNLRB2*) and the meiosis/spermiogenesis associated 1 (*MEIG1*), were the principal cells with higher levels of PSG expression (Fig. 4C; Table S5). Those “ciliated cells” were not anatomically identified in the cerebellum (Carter et al. 2018), but their associated cilium markers were found to be expressed at the site of the cerebellar granule cells (Lein et al. 2007). These cells may, therefore, be a subtype of granule neurons involved in cerebellar function. The PSG expressed in these ciliated cells code for the tubulin tyrosine ligase like 6 (*TTLL6*), *TOP3A*, the dynein cytoplasmic 2 light intermediate chain 1 (*DYNC2LI1*) and the lebercilin (*LCA5*) coding for a component of the axoneme of ciliated cells. Some of these PSG are also involved in human brain diseases such as microcephaly, macrocephaly, and Joubert syndrome (Fig. 4D and see below).

Finally, we assessed the potential association with brain functions, by extracting 19,244 brain imaging results from 315 fMRI-BOLD studies (T and Z score maps; see Table S6 for the complete list) from NeuroVault (Gorgolewski et al. 2015) and comparing the spatial patterns observed with the patterns of gene expression in the Allen Brain Atlas (Hawrylycz et al. 2012; Gorgolewski et al. 2014). The correlation between brain activity and PSG expression was stronger in subcortical structures than in the cortex (Wilcoxon rc=0.14, *p*=2.5×10^-248^). The brain activity maps that correlate with the expression pattern of the PSG (see Table S7 for details) were enriched in social tasks (empathy, emotion recognition, theory of mind, language; Fisher’s exact test *p*=2.9×10^-20^, OR=1.72, CI_95%_=[1.53, 1.93]). We also observed this enrichment for expression pattern of the SCG (Fisher’s exact test *p*=1.2×10^-12^, OR=1.16, CI_95%_=[1.11, 1.22]), however there were significantly less correlated than those of PSG (Fisher’s exact test *p*=0.0004, OR=0.83, CI_95%_=[0.75, 0.92]).

### Positively selected genes and their relationship to brain disorders

Our systematic analysis revealed that SCG were more associated with brain diseases or traits than PSG (Fig. 3B), particularly for intellectual disability (*p*=8.13×10^-6^, OR=0.34 CI_95%_=[0.21, 0.56], Bonferroni-corrected) and autism (*p*=0.0005, OR=0.26, CI_95%_=[0.11, 0.59], Bonferroni-corrected). We also identified 42 high PSG associated (based on OMIM and HPO data) with several human diseases or conditions, such as micro/macrocephaly, autism, or dyslexia (Table S4).

A comparison of humans and chimpanzees with our common primate ancestor revealed several protein-coding genes associated with micro/macrocephaly with different patterns of evolution in humans and chimpanzees (Fig. 5). Some genes displayed a divergence specifically in the hominin lineage (*AHI1, ASXL1, BRCA1, CSPP1, DAG1, FAM111A, FAM149B1, GRIP1, NHEJ1, QDPR, RNF135, RNF168, SLX4, TCTN1, TMEM70, TMEM260*, and *TOP3A*) or in the chimpanzee (*ALKBH8, ARHGAP31, ATRIP, CPT2, CTC1, HDAC6, HEXB, KIF2A, MKKS, MRPS22, RFT1, TBX6*, and *WWOX*).

**Figure 5.**
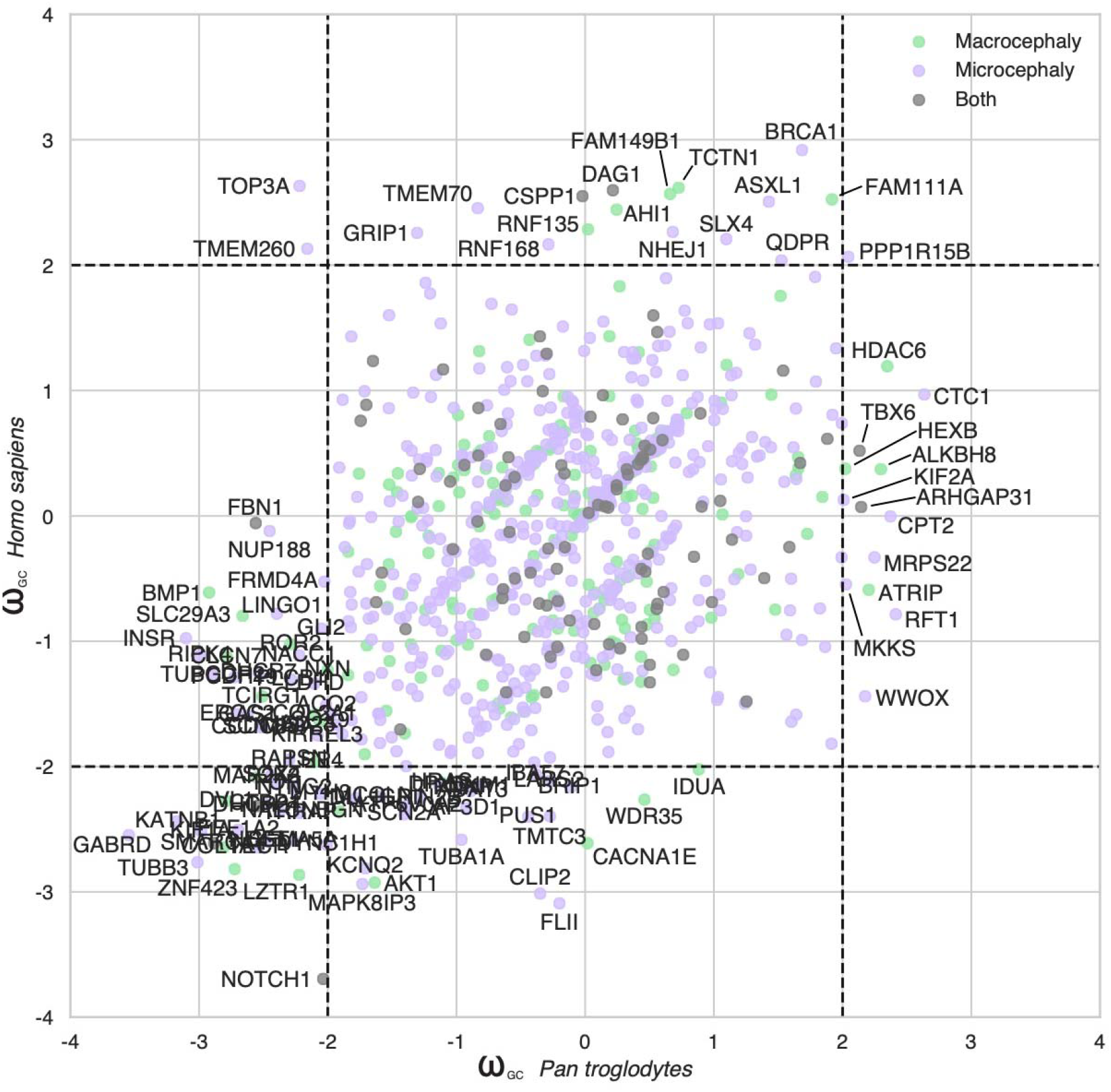
Evolution of the protein-coding genes associated with micro- or macrocephaly in humans. Scatter plots comparing ω_GC12_ between *Homo sapiens* and *Pan troglodytes* for the microcephaly- and macrocephaly-associated genes.

We also identified PSG associated with communication disorders, such as autism (*CNTNAP4, AHI1, FAN1, SNTG2*, and *GRIP1*) and dyslexia (*KIAA0319*). These genes diverged from the common primate ancestor only in the hominin lineage and were under strong selective constraint in all other taxa (Fig. 6A and 6B). They all have roles relating to neuronal connectivity (neuronal migration and synaptogenesis) and, within the human brain, were more specifically expressed in the cerebellum, except for *GRIP1*, which was expressed almost exclusively in the cortex.

**Figure 6.**
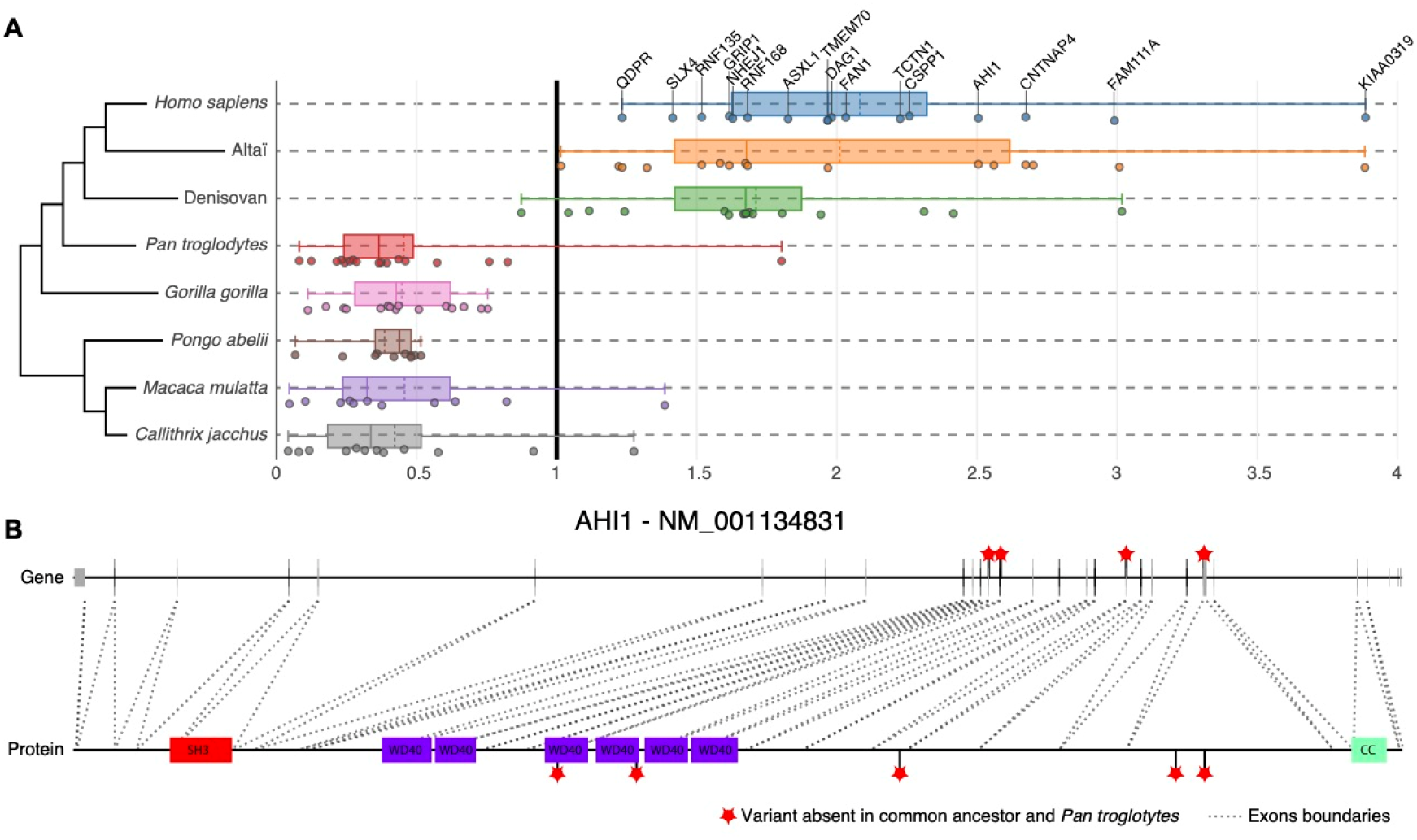
Examples of brain disorder-associated protein-coding genes displaying specific divergence in hominins during primate evolution. (A) Representation of 16 genes with *d*_N_/*d*_S_ >1 in *Homo sapiens* and archaic hominins but *d*_N_/*d*_S_ <1 for other primates. (B) Representation of hominin-specific non-synonymous variants of the *AHI1* gene, showing the correspondence with the protein (dot lines indicate exons); note how two variants lie within the WP40 functional domains. Red stars indicate variants (CADD>5) relative to the ancestor present in *Homo sapiens*, Neanderthals, and Denisovans, but not in *Pan troglodytes*. WP40: WD40 repeat; SH3: SRC Homology 3; CC: Coiled-coils.

The dyslexia susceptibility gene *KIAA0319*, encoding a protein involved in axon growth inhibition (Paracchini et al. 2006; Franquinho et al. 2017), is one of the PSG under the strongest positive selection in humans relative to the common primate ancestor (raw *d*_N_/*d*_S_ =3.9; 9 non-synonymous vs. 1 synonymous mutations in *Homo sapiens* compared to the common primate ancestor). The role of *KIAA0319* in dyslexia remains a matter of debate, but its rapid evolution in the hominoid lineage warrants further genetic and functional studies.

Finally, several PSG display very high levels of positive selection in *Homo sapiens*, but their functions or association with disease remain unknown. For example, the zinc finger protein ZNF491 (raw *d*_N_/*d*_S_ =4.7; 14 non-synonymous vs. 1 synonymous mutation in *Homo sapiens* compared to the common primate ancestor) is specifically expressed in the cerebellum and is structurally similar to a chromatin remodeling factor, but its biological role remains to be determined. Another example is the *CCP110* gene, encoding a centrosomal protein resembling ASPM, but not associated with a disease. Its function suggests that this PSG would be a compelling candidate for involvement in microcephaly in humans. A complete list of the brain SCG and PSG is available in Table S4 and on the companion website.

## Discussion

### Positively selected genes and brain size in primates

Several protein-coding genes are thought to have played a significant role in the increase in brain size in humans. Some of these genes, such as *ARHGAP11B, SRGAP2C*, and *NOTCH2NLA* (Suzuki et al. 2018), are specific to humans, having recently been duplicated (Dennis and Eichler 2016). Other studies have suggested that a high degree of positive selection in genes involved in micro/macrocephaly may have contributed to the substantial change in brain size during primate evolution (Dorus et al. 2004; Hayward 2004). Several of these genes, such as *ASPM* (Mekel-Bobrov et al. 2005) and *MCPH1* (Evans et al. 2005), seem to have evolved more rapidly in humans. However, the adaptive nature of the evolution of these genes has been called into question (Yu et al. 2007), and neither of these two genes were on the PSG list in our analysis (their raw *d*_N_/*d*_S_ value are below 0.8).

Conversely, our systematic detection approach identified the genes under the strongest positive selection in humans for micro/macrocephaly, the top 10 such genes being *AHSG, ASXL1, BRCA1, CSPP1, DAG1, FAM111A, FAM149B1, RNF168, TMEM70* and *TOP3A*. This list of PSG associated with micro/macrocephaly in humans can be used to select the best candidate human-specific gene/variants for further genetic and functional analyses, to improve estimates of their contribution to the emergence of anatomic difference between humans and other primates.

As previously shown, our systematic analysis confirms that the major susceptibility gene for breast cancer *BRCA1* is under strong positive selection (Lou et al. 2014). BRCA1 is a DNA damage response protein that repairs double-strand breaks in DNA. Heterozygous *BRCA1* mutations increase the risk of breast cancer, but can also cause neuronal migration defects (Eccles et al. 2005). In sporadic cases, homozygous *BRCA1* mutations lead to Fanconi anemia with microcephaly (Mehmet et al. 2016). Several other DNA damage response proteins (Arcas et al. 2014), which are binding partners of BRCA1 such as SLX4, TOP3A, RNF168 and MCPH1 are also associated with microcephaly. How *BRCA1* mutations cause microcephaly in humans remains largely unknown. However, in the mouse, *Brca1* mutations strongly reduce the size of the cerebral cortex by affecting the cellular polarity of neural progenitors and preventing the apoptosis of early cortical neuron progenitors (Jn and Wb 2009; Pao et al. 2014). Upper-most cortical layers are not reduced upon *Brca1* ablation in mice, and this is consistent with the low levels of apoptosis found in late progenitors and the neurons derived from there. Our analysis of the single-cell RNA-seq data from the human cortex indicates that excitatory neurons from layers 3 and 5 express PSG more than expected, including *BRCA1* and several of its binding partners associated with DNA damage response and microcephaly such as *TOP3A, RNF168*, and *NHEJ1*. Further analyses on the role of these genes, which are currently known for their DNA damage response, might shed some light on primate brain evolution.

In addition to brain size, some of the micro/macrocephaly PSG genes may have contributed to differences in other morphological features, such as skeleton development. For example, the PSG *FAM111A* (raw *d*_N_/*d*_S_ =2.99; 7 non-synonymous vs. 1 synonymous mutations in *Homo sapiens* compared to the common primate ancestor) and *ASXL1* (raw *d*_N_/*d*_S_ =1.83; 12 non-synonymous vs. 3 synonymous mutations in *Homo sapiens* compared to the common primate ancestor) are associated with macrocephaly and microcephaly, respectively. Patients with dominant mutations of *FAM111A* are diagnosed with Kenny-Caffey syndrome (KCS). They display impaired skeletal development, with small dense bones, short stature, primary hypoparathyroidism with hypocalcemia, and a prominent forehead (Unger et al. 2013). FAM111A is a binding partner of BRCA1 and plays a role in DNA damage response, but this protein seems to be also crucial to a pathway governing parathyroid hormone production, calcium homeostasis, and skeletal development and growth. By contrast, patients with dominant mutations of *ASXL1* are diagnosed with Bohring-Opitz syndrome, a malformation syndrome characterized by severe intrauterine growth retardation, intellectual disability, trigonocephaly, hirsutism, and flexion of the elbows and wrists with a deviation of the wrists and metacarpophalangeal joints (Hoischen et al. 2011). *ASXL1* encodes a chromatin protein required to maintain both the activation and silencing of homeotic genes.

Three genes (*AHI1, CSPP1*, and *TCTN1*) in the top 10 of the PSG associated with human brain diseases, with raw *d*_N_/*d*_S_ >2, are required for both cortical and cerebellar development in humans. They are also associated with Joubert syndrome, a recessive disease characterized by agenesis of the cerebellar vermis and difficulties coordinating movements. *AHI1* is a positive modulator of classical WNT/ciliary signaling. *CSPP1* is involved in cell cycle-dependent microtubule organization, and *TCTN1* is a regulator of Hedgehog during development.

*AHI1* was previously identified as a gene subject to positive selection during the evolution of the human lineage (Ferland et al. 2004; Gould and Walter 2004), but, to our knowledge, neither *CSPP1* nor *TCTN1* has previously been described as a diverging during primate evolution. It has been suggested that the accelerated evolution of *AHI1* required for ciliogenesis and axonal growth may have played a role in the development of unique motor capabilities, such as bipedalism, in humans (Hayward 2004). Our findings provide further support for the accelerated evolution of a set of genes associated with ciliogenesis.

### The possible link between a change in the genetic makeup of the cerebellum and the evolution of human cognition

The emergence of a large cortex was undoubtedly an essential step for human cognition, but other parts of the brain, such as the cerebellum, may also have made significant contributions to both motricity and cognition. In this study, we showed that the protein-coding genes expressed in the cerebellum were among the most conserved in humans. However, we also identified a set of PSG with relatively strong expression in the cerebellum or for which mutations affected the cerebellar function. As discussed above, several PSG are associated with Joubert syndrome, including *AHI1, CSPP1*, and *TCTN1*, and are essential for cerebellar development. Furthermore, the PSG expressed in the brain and under the highest positive selection include *CNTNAP4, FAN1, SNTG2*, and *KIAA0319*, which also display high levels of expression in the cerebellum and have been associated with communication disorders, such as autism and dyslexia. Finally, the choroid plexus expressed more PSG than expected and is known to play the role of a paracrine gland to produce the retinoic acid necessary for cerebellum development (Yamamoto et al. 1996).

In humans, the cerebellum is associated with higher cognitive functions, such as visuospatial skills, the planning of complex movements, procedural learning, attention switching, and sensory discrimination (Koziol et al. 2012). It plays a crucial role in temporal processing (Rao et al. 2001) and the anticipation and control of behavior through both implicit and explicit mechanisms (Koziol et al. 2012). A change in the genetic makeup of the cerebellum would, therefore, be expected to have been of great advantage for the emergence of the specific features of human cognition.

Despite this possible link between the cerebellum and the emergence of human cognition, much less attention has been paid to this part of the brain than to the cortex, on which most of the functional studies investigating the role of human-specific genes/variants have focused. For example, *SRGAP2C* expression is almost exclusively restricted to the cerebellum in humans, but the ectopic expression of this gene has been studied in mouse cortex (Charrier et al. 2012; Dennis et al. 2012), in which it triggers human-like neuronal characteristics, such as an increase in dendritic spine length and density. We thus suggest that an exploration of human genes/variants specifically associated with the development and functioning of the cerebellum might shed new light on the evolution of human cognition.

### Limitations

The present results have potential limits in their interpretations. Sources of error in the alignments (e.g., false orthologous, segmental duplications, errors in ancestral sequence reconstruction) are still possible and can result in inflated *d*_N_/*d*_S_. The *d*_N_/*d*_S_ method is not suited for comparing very closely related species and therefore, differences between *Neanderthal, Denisovan*, and *Homo sapiens* must be taken with care. Moreover, methods to estimate the evolution of proteins are expected to give downwardly biased estimates (Eyre-Walker and Keightley 2009). However, our GC12 normalization has already proved to correct for most of those biases in systematic analyses (Kapheim et al. 2015), and our raw *d*N/*d*S values highly correlate with other independent studies on primates (Biswas et al. 2016; Nielsen et al. 2005). Moreover, for the enrichment analyses, we used bootstrapping techniques to better control for potential biases induced by differences in GC content and gene length, especially for genes implicated in brain disorders (Zylka et al. 2015). Finally, our data are openly available on the companion website and allow to check at the variant level which amino acids changed.

### Perspectives

Our systematic analysis of protein sequence diversity confirmed that protein-coding genes relating to brain function are among the most highly conserved in the human genome. The set of PSG identified here may have played specific roles in the evolution of human cognition, by modulating brain size, neuronal migration, and synaptic physiology, but further genetic — including detailed analyses of all species branches— and functional studies would shed new light on the role of these genes. Beyond the brain, this resource will also be useful for estimating the evolutionary pressure acting on genes related to other biological pathways, particularly those displaying signs of positive selection during primate evolution, such as the reproductive and immune systems.

## Materials and Methods

### Genetic sequences

#### Alignments with the reference genome

We collected sequences and reconstructed sequence alignments with the reference human genome version hg19 (release 19, GRCh37.p13). For the primate common ancestor sequence, we used the Ensembl 6-way Enredo-Pecan-Ortheus (EPO) (Paten et al. 2008) multiple alignments v71, related to *Homo sapiens* (hg19), chimpanzee (panTro4), gorilla (gorGor3), orangutan (ponAbe2), rhesus macaque (rheMac3), and marmoset (calJac3). For the two ancestral hominins, Altai, and Denisovan, we integrated variants detected by Castellano and colleagues (Castellano et al. 2014) into the standard hg19 sequence (http://cdna.eva.mpg.de/neandertal/, date of access 2014-07-03). Finally, we used the whole-genome alignment of all the primates used in the 6-EPO from the UCSC website (http://hgdownload.soe.ucsc.edu/downloads.html, access online: August 13, 2015). All the PSG had their protein sequence deduced from our analysis compared manually to the one in the protein database. All variants matched and we did not find any alignment artifact. The core annotations used for our study were not available for the GRCh38 version of the human genome when we started this project. Since one of the biggest improvements in GRCh38 is the annotation of the centromere regions (Guo et al. 2017), a switch from GRCh37 to GRCh38 would not affect our conclusions. Moreover, regarding the coding regions of the human genome, the number of nonsynonymous detected by GRCh38 (N=22,796 SNVs) is very similar to GRCh37’s (N=22,622 SNVs; see Table 3 in Guo et al. 2017).

#### VCF annotation

We combined the VCF file from Castellano and colleagues (Castellano et al. 2014) with the VCF files generated from the ancestor and primate sequence alignments. The global VCF was annotated with ANNOVAR (Wang et al. 2010) (version of June 2015), using the following databases: refGene, cytoBand, genomicSuperDups, esp6500siv2_all, 1000g2014oct_all, 1000g2014oct_afr, 1000g2014oct_eas, 1000g2014oct_eur, avsnp142, ljb26_all, gerp++elem, popfreq_max, exac03_all, exac03_afr, exac03_amr, exac03_eas, exac03_fin, exac03_nfe, exac03_oth, exac03_sas. We also used the ClinVar database (https://ncbi.nlm.nih.gov/clinvar/, date of access 2016-02-03).

### ω_GC12_ calculation

Once all the alignments had been collected, we extracted the consensus coding sequences (CCDS) of all protein-coding genes referenced in Ensembl BioMart Grc37, according to the HGNC (date of access 05/05/2015) and NCBI Consensus CDS protein set (date of access 2015-08-10). We calculated the number of non-synonymous mutations N, the number of synonymous mutations S, the ratio of the number of non-synonymous mutations per non-synonymous site dN, the number of synonymous mutations per synonymous site dS, and their ratio *d*_N_/*d*_S_ — also called □—between all taxa and the ancestor, using the yn00 algorithm implemented in PamL software (Yang 2007). We avoided infinite and null results by calculating a corrected version of *d*_N_/*d*_S_. If S was null, we set its value to one to avoid having zero as the numerator. The obtained values were validated through the replication of a recent systematic estimation of *d*_N_/*d*_S_ between *Homo sapiens* and two great apes (Biswas et al. 2016) (*Pan troglodytes* and *Pongo abelii;* Pearson’s r>0.8, p<0.0001; see Fig. S2). Finally, we obtained our ω_GC12_ value by correcting for the GC12 content of the genes with a generalized linear model and by calculating a *Z*-score for each taxon (Kapheim et al. 2015). GC content has been associated with biases in mutation rates, particularly in primates (Galtier et al. 2009) and humans (Kostka et al. 2012). We retained only the 11667 genes with 1:1 orthologs in primates (extracted for GRCh37.p13 with Ensembl BioMart, access online: February 27, 2017).

### Gene sets

We used different gene sets, starting at the tissue level and then focusing on the brain and key pathways. For body tissues, we used Illumina Body Map 2.0 RNA-seq data, corresponding to 16 human tissue types: adrenal, adipose, brain, breast, colon, heart, kidney, liver, lung, lymph, ovary, prostate, skeletal muscle, testes, thyroid, and white blood cells (for more information: https://personal.broadinstitute.org/mgarber/bodymap_schroth.pdf; data preprocessed with Cufflinks, accessed May 5, 2015 at http://cureffi.org). We also used the microarray dataset of Su and colleagues (Su et al. 2004) (Human U133A/GNF1H Gene Atlas, accessed May 4, 2015 at http://biogps.org). Finally, we also replicated our results with recent RNA-seq data from the GTEx Consortium (2015; https://www.gtexportal.org/home/).

For the brain, we used the dataset of Su and colleagues and the Human Protein Atlas data (accessed November 7, 2017 at https://www.proteinatlas.org). For analysis of the biological pathways associated with the brain, we used KEGG (accessed February 25, 2015, at http://www.genome.jp/kegg/), synaptic genes curated by the group of Danielle Posthuma at Vrije Universiteit (accessed September 1, 2014, at https://ctg.cncr.nl/software/genesets), and mass spectrometry data from Loh and colleagues (Loh 2016). Finally, for the diseases associated with the brain, we combined gene sets generated from Human Phenotype Ontology (accessed August 14, 2020, at http://human-phenotype-ontology.github.io) including OMIM annotation (https://omim.org), and curated lists: the 65 risk genes proposed by Sanders and colleagues (Sanders et al. 2015) (TADA), the candidate genes for autism spectrum disorders from SFARI (accessed July 17, 2015 at https://gene.sfari.org), the Developmental Brain Disorder or DBD (accessed July 12, 2016 at https://geisingeradmi.org/care-innovation/studies/dbd-genes/), and Cancer Census (accessed November 24, 2016 at cancer.sanger.ac.uk/census) data. Note that the combination of HPO & OMIM is the most exhaustive, making it possible to avoid missing potential candidate genes, but this combination does not identify specific associations.

SynGO was generously provided by Matthijs Verhage (access date: January 11, 2019). This ontology is a consistent, evidence-based annotation of synaptic gene products developed by the SynGO consortium (2015-2017) in collaboration with the GO-consortium. It extends the existing Gene Ontology (GO) of the synapse and follows the same dichotomy between biological processes (BP) and cellular components (CC).

For single-cell transcriptomics datasets, we identified the genes specifically highly expressed in each cell type, following the same strategy as used for the other RNA-seq datasets. The single-cell data for the developing human cortex were kindly provided by Maximilian Haeussler (available at https://cells.ucsc.edu; access date: October 30, 2018). The single-cell transcriptional atlas data for the developing murine cerebellum (Carter et al. 2018) were kindly provided by Robert A. Carter (access date: January 29, 2019). For each cell type, we combined expression values cross all available replicates, to guarantee a high signal-to-noise ratio. We then calculated the values for the associated genes in *Homo sapiens* according to the paralogous correspondence between humans and mice (Ensembl BioMart accessed on February 23, 2019).

### Gene nomenclature

We extracted all the EntrezId of the protein-coding genes for Grc37 from Ensembl BioMart. We used the HGNC database to recover their symbols. For the 46 unmapped genes, we searched the NCBI database manually for the official symbol.

### McDonald-Kreitman-test (MK), neutrality index (NI), and Direction of Selection (DoS)

We assessed the possible fixation of variants in the *Homo sapiens* population by first calculating the relative ratio of non-synonymous to synonymous polymorphism (pN/pS) from the 1000 Genomes VCF for all SNPs, for SNPs with a minor allele frequency (MAF) <1% and <5%. SNPs were annotated with ANNOVAR across 1000 Genomes Project (ALL+5 ethnicity groups), ESP6500 (ALL+2 ethnicity groups), ExAC (ALL+7 ethnicity groups), and CG46 (see http://annovar.openbioinformatics.org/en/latest/user-guide/filter/#popfreqmax-and-popfreqall-annotations_for_more_details). The polymorphism ratio (pN/pS) allowed us to takes into account the constraint on nonsynonymous sites and thus increase the power of detecting positive selection (Salvador-Martínez et al. 2018). We indeed normalized the divergence ratio (*d*_N_/*d*_S_) using the McDonald–Kreitman test i.e. calculating the neutrality index (NI) as the ratio of raw *p*_N_/*p*_S_ and *d*_N_/*d*_S_ values (McDonald and Kreitman 1991). We considered the PSG to be fixed in the population when NI < 1. We also confirmed with a new statistic for evolutionary measure: the Direction of Selection (DoS) = *D*_n_/(*D*_n_ + *D*_s_) – *P*_n_/(*P*_n_ + *P*_s_) (Stoletzki and Eyre-Walker 2011) that all divergent genes with NI<0 had a DoS < 0 (Fig. S8).

### NeuroVault analyses

We used the NeuroVault website (Gorgolewski et al. 2015) to collect 19,244 brain imaging results from fMRI-BOLD studies (*T* and *Z* score maps) and their correlation with the gene expression data (Gorgolewski et al. 2014) of the Allen Brain Atlas (Hawrylycz et al. 2012). The gene expression data of the Allen Brain atlas were normalized and projected into the MNI152 stereotactic space used by NeuroVault, using the spatial coordinates provided by the Allen Brain Institute. An inverse relationship between cortical and subcortical expression dominated the pattern of expression for many genes. We thus calculated the correlations for the cortex and subcortical structures separately.

### Allen Brain data

We downloaded the Allen Brain atlas microarray-based gene data and multiple cortical areas - Smart-seq from the Allen Brain website (accessed July, 2020 at http://www.brain-map.org). Microarray data were available for six adult brains; the right hemisphere was missing for three donors, so we considered only the left hemisphere for our analyses. For each donor, we averaged probes targeting the same gene and falling in the same brain area. We then subjected the data to log normalization and calculated *Z*-scores: across the 20787 genes for each brain region to obtain expression levels; across the 212 brain areas for each gene to obtain expression specificity. For genes with more than one probe, we averaged the normalized values over all probes available. The Smart-seq dataset followed a similar preprocessing and lead to expression level and specificity of 32165 genes across 363 cell types.

As a complementary dataset, we also used a mapping of the Allen Brain Atlas onto the 68 brain regions of the Freesurfer atlas (French and Paus 2015) (accessed April 4, 2017 at https://figshare.com/articles/A_FreeSurfer_view_of_the_cortical_transcriptome_generated_from_the_Allen_Human_Brain_Atlas/1439749). The expression and specificity measure were used for the 3D visualization in the companion website.

### Statistics

#### Enrichment analyses

We first calculated a two-way hierarchical clustering on the normalized *d*_N_/*d*_S_ values (**ω**_GC_) across the whole genome (see Fig. 1B; note: 11,667 genes were included in the analysis to ensure medium-quality coverage for *Homo sapiens*, Neanderthals, Denisovans, and *Pan troglodytes;* see Fig. S1). According to 30 clustering indices (Charrad et al. 2014), the best partitioning in terms of evolutionary pressure was into two clusters of genes: SCG (*N*=4825; in HS, mean=-0.88 median=-0.80 SD=0.69) and PSG (N=6842; in HS, mean=0.60 median=0.48 sd=0.63. For each cluster, we calculated the enrichment in biological functions in Cytoscape (Shannon et al. 2003) with the BINGO plugin (Maere et al. 2005). We used all 11,667 genes as the background. We eliminated redundancy, by first filtering out all the statistically significant Gene Ontology (GO) terms associated with fewer than 10 or more than 1000 genes, and then combining the remaining genes with the EnrichmentMap plugin (Merico et al. 2010). We used a *P*-value cutoff of 0.005, an FDR Q-value cutoff of 0.05, and a Jaccard coefficient of 0.5.

For the cell type-specific expression analysis (CSEA; 86), we used the CSEA method with the online tool http://genetics.wustl.edu/jdlab/csea-tool-2/. This method associates gene lists with brain expression profiles across cell types, regions, and time periods.

#### Wilcoxon and rank-biserial correlation

We investigated the extent to which each gene set was significantly more under positive or constraint selection than expected by chance, by performing Wilcoxon tests on the normalized *d*_N_/*d*_S_ values (ω_GC_) for the genes in the set against zero (the mean value for the genome). We quantified effect size by matched pairs rank-biserial correlation, as described by Kerby (Kerby 2014). Following non-parametric Wilcoxon signed-rank tests, the rank-biserial correlation was evaluated as the difference between the proportions of negative and positive ranks over the total sum of ranks:

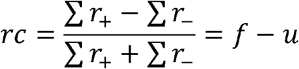

It corresponds to the difference between the proportion of observations consistent with the hypothesis (f) minus the proportion of observations contradicting the hypothesis (u), thus representing an effect size. Like other correlational measures, its value ranges from minus one to plus one, with a value of zero indicating no relationship. In our case, a negative rank-biserial correlation corresponds to a gene set in which more genes have negative *ω_GC_* values than positive values, revealing a degree of conservation greater than the mean for all genes (i.e., *ω_GC_* = 0). Conversely, a positive rank-biserial correlation corresponds to a gene set that is more under positive selection than expected by chance (i.e., taking randomly the same number of genes across the whole genome; correction for the potential biases for GC content and CDS length are done at the bootstrap level). All statistics relating to the Figures 1D, 2A, and 2B are summarized in the Table S3. All those relating to the Figures 4 are summarized in the Table S5.

#### Validation by resampling

We also used bootstrapping to correct for potential bias in the length of the coding sequence or the global specificity of gene expression (Tau, see the methods from Kryuchkova-Mostacci and Robinson-Rechavi in (Kryuchkova-Mostacci and Robinson-Rechavi 2016)). For each of the 10000 permutations, we randomly selected the same number of genes as for the sample of genes from the complete set of genes for which *d*_N_/*d*_S_ was not missing. We corrected for CCDS length and GC content by bootstrap resampling. We estimated significance, to determine whether the null hypothesis could be rejected, by calculating the number of bootstrap draws (*B*_i_) falling below and above the observed measurement (m). The corresponding empirical *p*-value was calculated as follows:

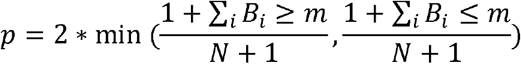

## Supporting information

Supplemental Table S1. Genes dN/dS.

Supplemental Table S2. GO Enrichment.

Supplemental Table S3. Statistics for all dNdS analyses.

Supplemental Table S4. List of brain SCG and PSG.

Supplemental Table S5. Statistics for scRNAseq.

Supplemental Table S6. List of the NeuroVault studies.

Supplemental Table S7. Social vs. Non-social NeuroVault analysis.

## Data access

All the data and code supporting the findings of this study are available from our resource website https://genevo.pasteur.fr and as Supplemental Material.

## Competing interest statement

The authors declare no competing interests.

## Acknowledgments

We thank J-P. Changeux, L. Quintana-Murci, E. Patin, G. Laval, B. Arcangioli, D. DiGregorio, L. Bally-Cuif, A. Chedotal, C. Berthelot, H. Roest Crollius, and V. Warrier for advice and comments, and the members of the Human Genetics and Cognitive Functions laboratory for helpful discussions. We also thank C. Gorgolewski, R. Carter, M. Haeussler, M. Verhage, and the SynGO consortium for providing key datasets without which this work would not have been possible. This work was supported by the Institut Pasteur; *Centre National de la Recherche Scientifique*; Paris Diderot University; the *Fondation pour la Recherche Médicale* [DBI20141231310]; the Human Brain Project; the Cognacq-Jay Foundation; the Bettencourt-Schueller Foundation; the *Agence Nationale de la Recherche* (ANR) [SynPathy]; and the Innovative Medicines Initiative 2 [no. 777394]. This research was also supported by the Laboratory of Excellence GENMED (Medical Genomics) grant no. ANR-10-LABX-0013, Bio-Psy, and by the INCEPTION program ANR-16-CONV-0005, all managed by the ANR part of the Investments for the Future program. G.D. is funded by the Institute for Data Valorization (IVADO), Montreal, and the Fonds de recherche du Québec (FRQ). The funders had no role in study design, data collection, and analysis, the decision to publish, or preparation of the manuscript.

## Author contributions

G.D. and T.B. devised the project and came up with the main conceptual ideas. G.D. developed the methods, performed the analyses, and designed the figures. G.D. and T.B. discussed the results and wrote the manuscript. S.M. developed the companion website.

## Notes

http://genevo.pasteur.fr

### Competing Interest Statement

The authors have declared no competing interest.

### Summary of Updates

Another round of edits with clarification about the dN/dS measure. Addition of new single-cell RNA seq data with convergence on a pathway containing BRCA1.

