## Supplemental methods and figures for "Systematic detection of brain protein-coding genes under positive selection during primate evolution and their roles in cognition"

Short title: **Evolution of brain protein-coding genes in humans**

Guillaume Dumas^a,b^, Simon Malesys^a^, and Thomas Bourgeron^a^

**^a^** Human Genetics and Cognitive Functions, Institut Pasteur, UMR3571 CNRS, Université de Paris, Paris, (75015) France

**^b^** Department of Psychiatry, Université de Montreal, CHU Ste Justine Hospital, Montreal, QC, Canada.

Supplemental Methods

**Sequence coverage**

We computed the overall analyses for three levels of sequence coverage: low, medium, and high with respectively 20%, 50%, or 80% of coverage of the consensus coding sequence (CCDS). For the core analyses of the paper, we focused on the genes with medium coverage in *Homo sapiens* (hg19), Neanderthal (altai), Denisovan, and *Pan troglodytes* (panTro4).

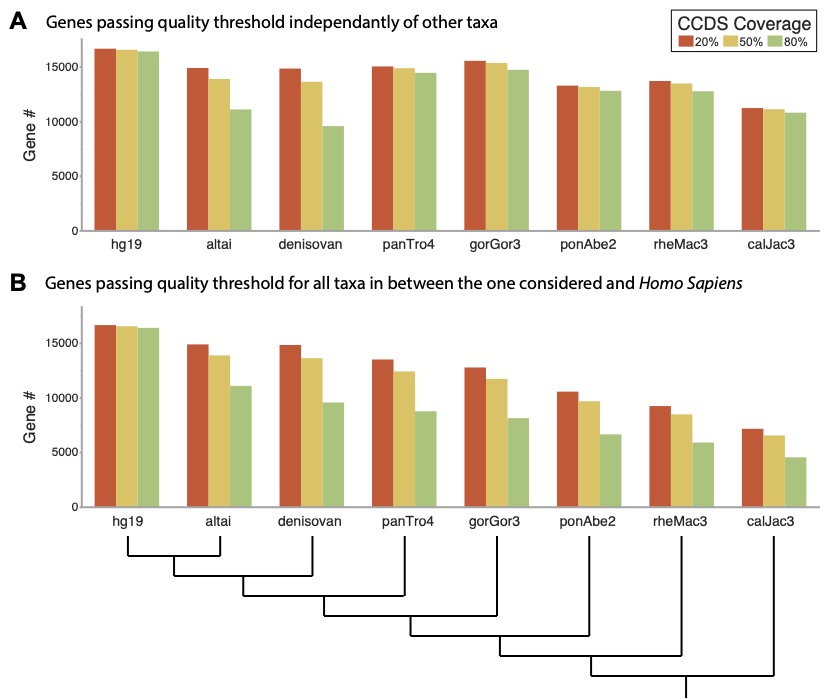

**Supplemental Fig S1.** **Quality check for the three levels of sequence coverage: low, medium, and high corresponding respectively to a coverage of the consensus coding sequence of at least 20%, 50%, and 80%. The number of protein-coding genes is calculated for either each taxon alone (A) or with all the taxa between *Homo sapiens* and the one considered (B).**

**Validation of dN/dS values**

We compared our systematic calculation of dN, dS, and dN/dS ratio with data recently published for primates (Biswas et al. 2016). Biswas and colleagues (2016) quantified the changes between *Homo sapiens* and two great apes: *Pan troglodytes* and *Pongo abelii*. Figure S2 summarizes all Pearson’s correlations between those published results and ours.

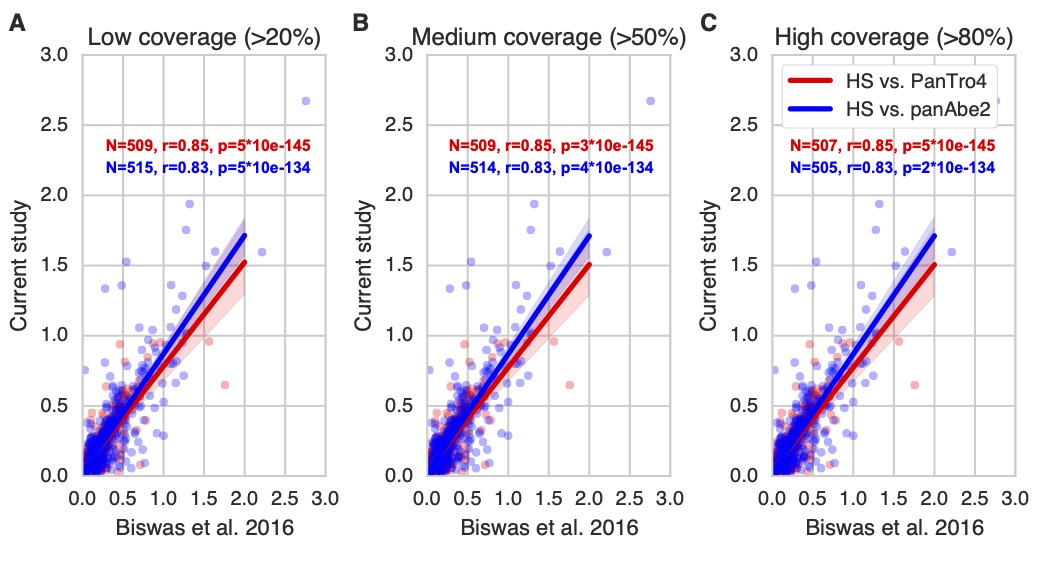

**Supplemental Fig S2. Comparison between dN/dS values obtained in the current study and the recently published paper by Biswas and colleagues (2016). Panels (A), (B), and (C) are respectively for low, medium, and high coverage of the CCDS used in the current study.**

**Highly expressed genes vs. tissue specific genes**

For identifying genes specifically transcribed in a given tissue, we computed a Z-score across all the tissues. We also computed global specificity index Tau for each gene following the methods from Kryuchkova-Mostacci and Robinson-Rechavi (2016) (Kryuchkova-Mostacci and Robinson-Rechavi 2016). Figure S3 illustrates the relationship between specificity and level of expression for the human brain. Note how genes with high expression levels are not necessarily specific to this tissue. Our study used the genes with expression specificity higher than 2 standard deviation (SD) as specific to the tissue.

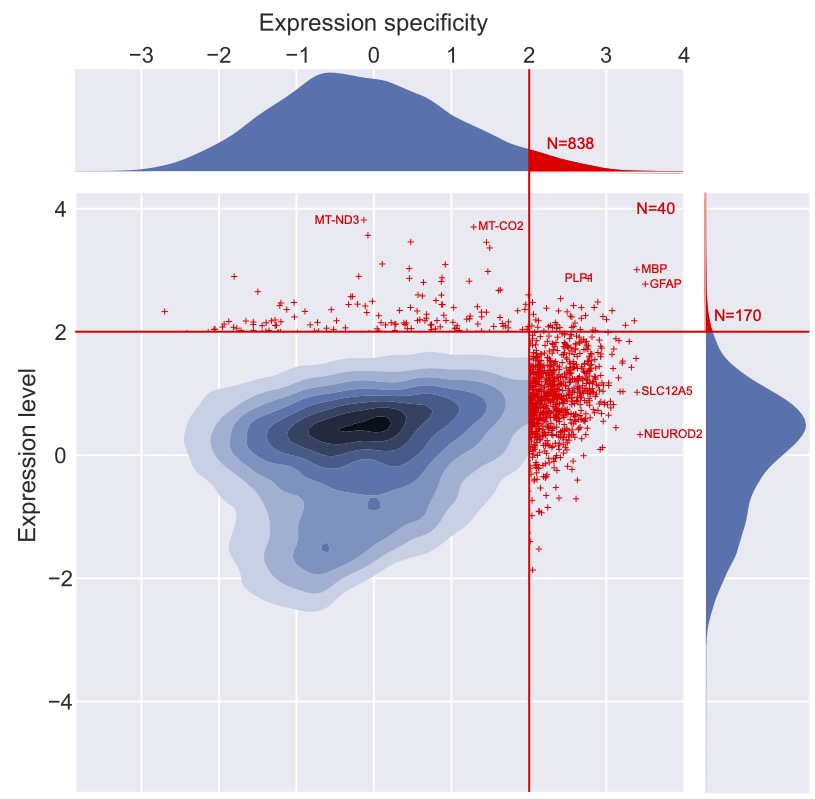

**Supplemental Fig S3. Illustration of the paper's framework to select genes specific to a given tissue, here the brain. Notice how the level of expression is not necessarily high.**

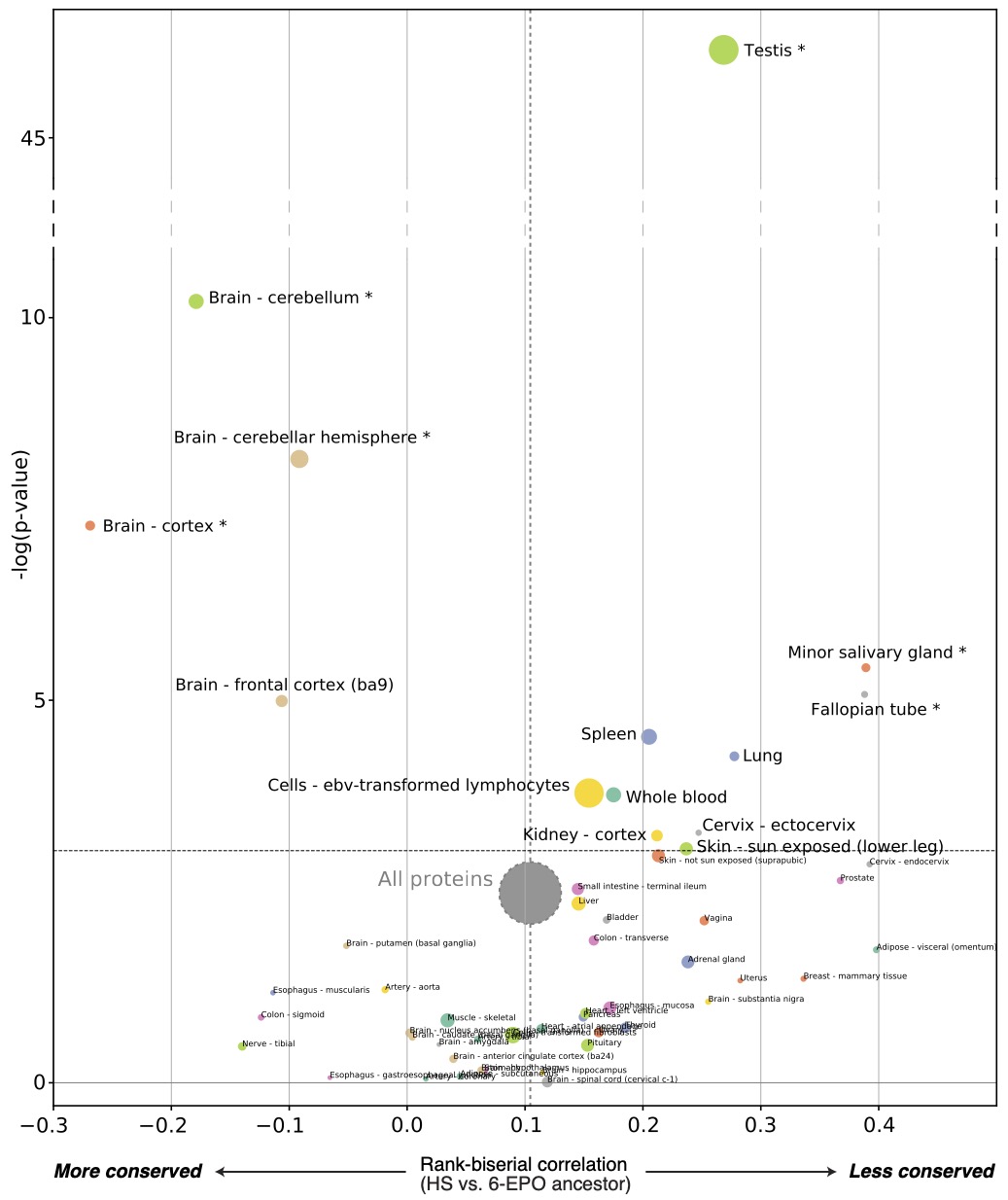

**Supplemental Fig S4. Replication of the BodyMap panel (d) in Figure 1 with the Genotype-Tissue Expression (GTEx) project, an RNAseq dataset for the scientific community to study the relationship between genetic variation and gene expression in human tissues (The GTEx Consortium 2015). The dashed horizontal line indicates the threshold for significance after Bonferroni correction. Stars indicate sets of genes for which statistical significance was achieved for multiple comparisons with bootstrap correction.**

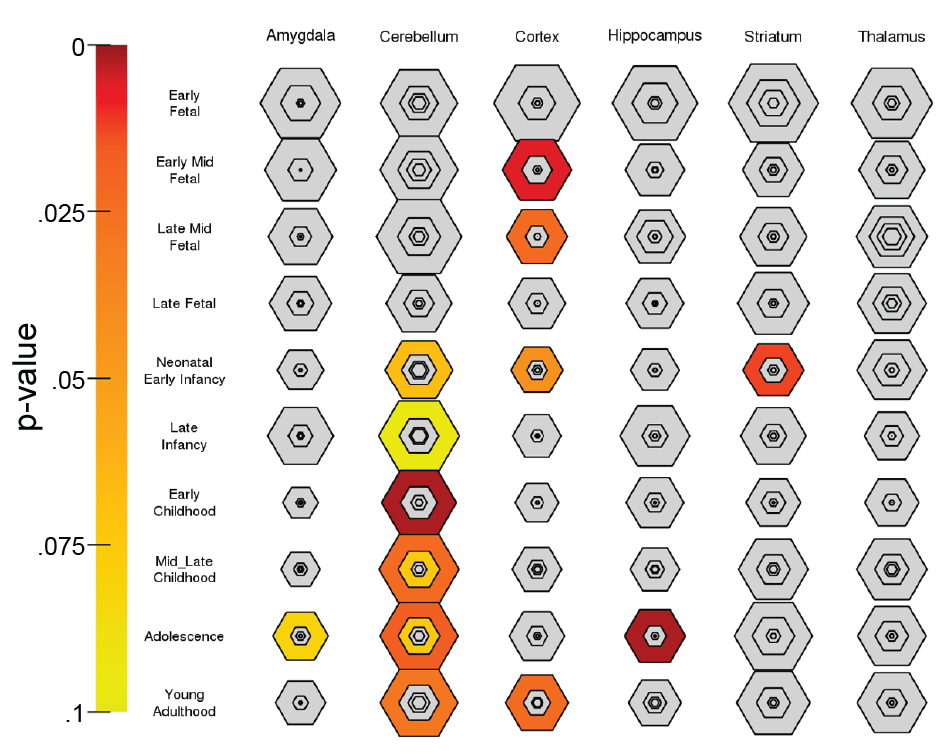

**Supplemental Fig S5. Specific Expression Analysis (SEA) across brain regions and development of the most conserved genes in *Homo Sapiens*. Varying stringencies for enrichment in Specificity Index thresholds (pSI) are represented by the size of the hexagons going from least specific lists (outer hexagons) to most specific (center).**

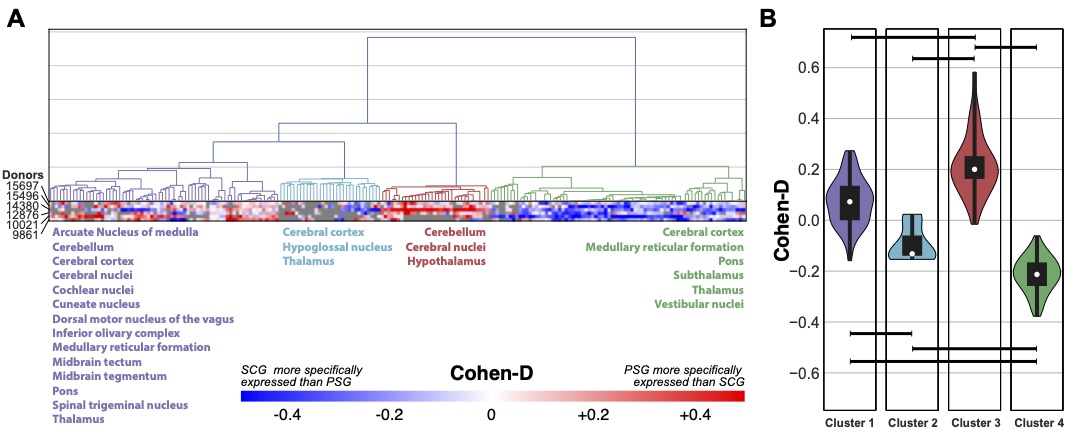

**Supplemental Fig S6. Differential expression signature of evolution across subgroups of cortical and subcortical human brain structures. (A) Hierarchical clustering on the different parts of the brain from the six donors in the Allen Brain atlas according to the differences in expression specificity of the PSG and SCG. Brain regions with similar evolutionary signature are indicated in the same colour under each of the four identified clusters. PSG are more specifically expressed in the brain regions from cluster 3. Notice that it does not contradict that cerebellum is conserved on average since only PSG and SCG are under consideration. This would support that even tissue expressing many conserved genes can also express more than expected genes under extreme positive selection. (B) Violin plot indicating the effect sizes of those differences in expression specificity averaged across the six brain donors for each brain structure in each of the 4 clusters (Kruskal-Wallis test W=134.8, p<0.001; Horizontal black bars indicate post hoc comparison with Mann-Whitney passing Bonferroni correction).**

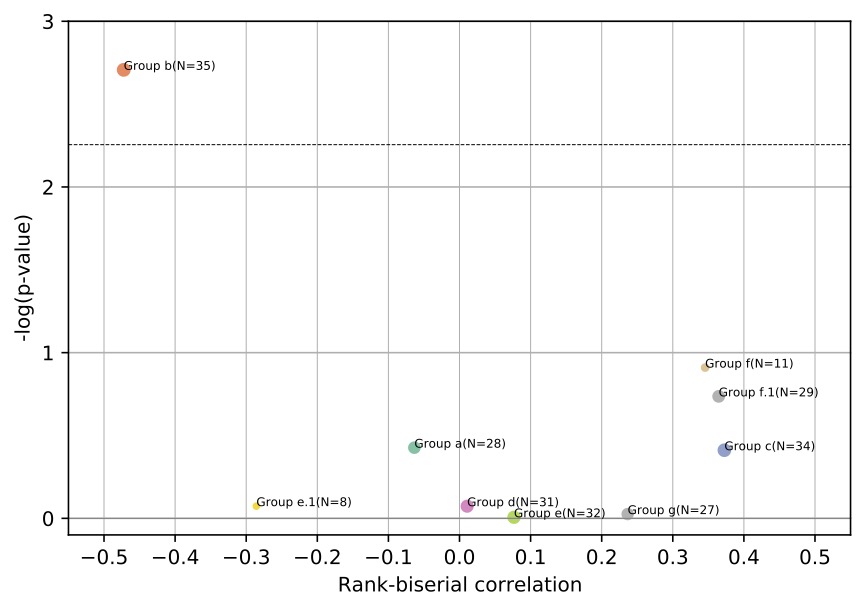

**Supplemental Fig S7. Evolution of protein-coding genes related to human cerebral organoids. Groups: a. Cell cycle, Forebrain development; b. Neuron diff. & projection; c. Cell adhesion, Vesicle transport; d. Neurogenesis, Cell migration; e. Cell adhesion, Cell morphogenesis (e.1 PC2 rg cor); F: Cell cycle, Mitosis (f.1. PC4 anti); G: Neurogenesis, Cell morphogenesis. For more details, see Camp et al. (2015). The dashed horizontal line indicates the threshold for significance after Bonferroni correction. Stars indicate sets of genes for which statistical significance was achieved for multiple comparisons with bootstrap correction.**

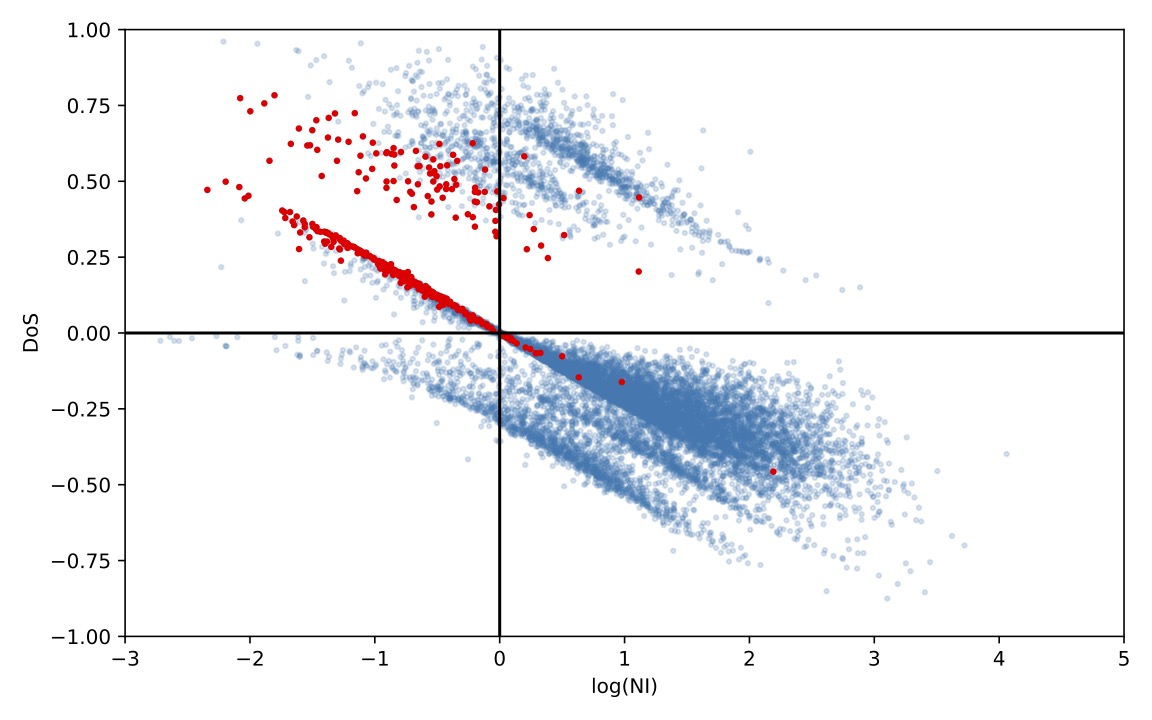

**Supplemental Fig S8. Comparison of NI and DoS. Each dot represents a gene. Red dots indicate PSG. Notice the absence of PSG with DoS < 0 and NI < 1.**
