## Supplemental Table S7. Social vs. Non-social NeuroVault analysis. for "Systematic detection of brain protein-coding genes under positive selection during primate evolution and their roles in cognition"

|  | **Studies with brain activity maps correlated  with pattern of expression of the divergent genes** | | |
| --- | --- | --- | --- |
|  | **Cortex** | **Subcortical structures** | **Both** |
| **All studies**  N=19244  Social: 7184 (37.33%) | N=1309  Social: 655 (50.04%) | N=1298  Social: 503 (38.75%) | N=95  Social: 47 (49.47%) |
| **Studies with  multiple subjects**  N=2569  Social: 319 (12.41%) | N=96  Social: 10 (10.41%) | N=228  Social: 33 (14.47%) | N=1  Social: 0 |

**Supplemental Table 7.** **Social vs. Non-social NeuroVault analysis.** Summary of the statistically significant correlations between the fMRI statistical maps from *Neurovault* and the expression pattern of PSG in either cortex or subcortical structure (bootstrap correcting for GC12 content and CDS length). The column “Both” refer to the number of maps both significantly correlated to cortex and subcortical structures. The grey cell indicates a statistical enrichment for social tasks (Fisher exact test, p=2.8e-20).
